# Quantifying Immune-Based Counterselection of Somatic Mutations

**DOI:** 10.1101/219576

**Authors:** Fan Yang, Dae-Kyum Kim, Hidewaki Nakagawa, Shuto Hayashi, Seiya Imoto, Lincoln Stein, Frederick P. Roth

## Abstract

It is now well established that somatic mutations in protein-coding regions can generate ‘neoantigens’, and that these can be recognized by the immune system and contribute to clearance of developing cancers. However, there is currently no model that can quantitatively predict the neoantigenic effect of any given somatic mutation. Here, we examined signatures of immune selection pressure on the distribution of somatic mutations. We quantified the extent to which somatic mutations are significantly depleted in peptides that are predicted to be displayed by major histocompatibility complex (MHC) class I proteins. We characterized the dependence of this depletion on expression level. We then examined whether immune selection pressure on somatic mutations changes depending on whether the patient had either one or two MHC-encoding alleles that can display the peptide. Our results indicate that MHC-encoding alleles are, in general, incompletely dominant, i.e., that having two copies of the display-enabling allele is more effective in displaying that peptide than having just one copy. More generally, a quantitative understanding of counter-selection of identifiable subclasses of neoantigenic somatic variation could guide immunotherapy or aid in developing personalized cancer vaccines.

## INTRODUCTION

In every human cell, proteins are constantly being degraded into component peptides, and a subset of this pool of peptides are displayed on MHC class I receptor proteins (encoded by human leukocyte antigen or HLA genes). As somatic mutations arise, some cause differences in MHC-displayed peptides, producing antigens that can be differentially recognized by T cells and lead the specific destruction of tumor cells by the immune system^1^. In addition to the production and display of ‘non-self’ peptides that can arise directly from mutation, genetic and epigenetic alterations can cause tumor cells to express many proteins more highly^2^. Together, these changes mean that cancer cells have an altered repertoire of proteins and therefore of tumor antigens.

Tumor antigens can be classified into two categories: tumor-associated self-antigens (which may be displayed by other normal cell types even if not displayed by the normal cell type from which the tumor was derived) and antigens derived from tumor-specific mutant proteins. The latter class of tumor-specific ‘neo-antigenic’ mutations are ideal targets for cancer immunotherapy, due to the fact that neo-antigens are less likely to be present in healthy cells/tissues and can potentially be recognized by the mature T-cell repertoire^3^. Also, it has been reported that neo-antigens are likely to be more immunogenic, presumably due to the T-cell maturation process in which T-cells capable of high-avidity recognition of self-antigens are eliminated^4^. Immuno-therapy approaches exploiting neo-antigenicity, however, have been hampered by the fact that every tumor possesses a unique set of mutations that must first be identified^5^. Moreover, individual patients can differ dramatically in their immune systems, based on HLA type and other allelic variation in immune genes, as well their unique repertoire of mature immune cells. Thus, personalized immuno-therapy could positively benefit the patient during cancer treatment^6,7,8^. After recognition, the process of tumor cell killing by T-cells may release more tumor neo-antigens in a potentially therapeutic virtuous cycle.

In principle, any coding mutations has the potential to generate mutant peptides that can be presented by MHC class I molecules and subsequently recognized by cytotoxic T cells. However, to bring this personalized treatment approach to tumor patients, a crucial challenge is determining the MHC-binding potential of non-self peptides that arise from somatic tumor mutations, and determining which among them are most likely to be potent neo-antigens in patients in different cancers with different HLA alleles that encode different MHC class I receptors.

To improve our understanding of neo-antigenicity in cancer, we conducted several analyses of somatic mutations and the ability of corresponding mutant peptides to be displayed by MHC class I receptors across different cancer types. More specifically, we quantified the impact of predicted antigenicity on the spectrum of tumor missense somatic mutations. We expected to find that somatic mutations would be less frequent in MHC-displayed peptides, presumably because of counter-selection of cells bearing these mutations by the immune system. Other groups have identified predicted-displayed mutations based on patient HLA-A genotypes^9^, without quantifying the extent of depletion of these mutations. Other work reported that predicted-MHC-displayed mutations were depleted in colorectal and clear cell renal cancer^10^. However, this phenomenon was not explored in detail, e.g., considering patient genotypes at all HLA loci or considering expression levels of the displayed peptide.

The project consisted of three main parts. First, we quantified the extent to which somatic mutations are significantly depleted in peptides that are predicted to be displayed by MHC class I proteins (without considering patient HLA type). We characterized the dependence of this depletion on the inferred expression level of each peptide. Second, we refined each of the preceding analyses by considering individual patient HLA alleles. Third, we extended this analysis by relating depletion of somatic mutations to the number of HLA alleles predicted to display peptides bearing that mutation. Thus, we quantitatively estimated the ‘neoantigenicity’ of different classes of somatic variants in individual patients.

## RESULTS

### Depletion of mutations within expressed predicted MHC-binding peptides

As somatic mutations arise, we should expect that more immunogenic mutations are more likely to be counter-selected due to clearance of the mutant cell by the immune system, and therefore depleted from observed tumor genomes. To formally test this hypothesis and to begin to quantify the expected depletion effect, we examined somatic cancer mutations in human cancer samples, beginning with whole genome/exome sequencing studies in the PCAWG database.

The immunogenicity of a protein-coding mutation depends in part on whether or not it yields a mutant peptide that is displayed by a MHC class I protein receptor. MHC class I binding peptides were therefore predicted using the NetMHC server^11,12^. In total, we examined 121,258 missense somatic mutations from 2,834 PCAWG patients for whom HLA type was estimated. Those mutations are distributed in more than 10,700 genes. Missense somatic mutations from PCAWG were separated into two groups: either falling within or outside of predicted MHC binding peptides. For an initial analysis, we modeled all MHC class I alleles for which display predictions were available as being present in each patient (we revisit this issue later).

Because a mutant protein must be expressed in order to yield a displayed peptide, we also examined the dependence of missense variant depletion on gene expression levels. More specifically, we analyzed the relationship between the missense mutation density within MHC binding peptides and the expression level of the corresponding protein using RNAseq data matched to the appropriate cancer type (see Materials and Methods). Then, for mutations within and outside of MHC binding peptides, we calculated the mutation density for five classes of peptide: those that were undetectably expressed and those in each of four gene expression quantiles (Materials and Methods).

As expected, we found that mutation density and expression level are negatively correlated, and that the average mutation density within MHC binding peptides is lower than that of MHC binding peptides for expressed peptides (Figure 1; ratio of mutation density within MHC-displayed peptides to that outside displayed peptides = 0.94; Fisher’s exact test, *P*-value < 2.2e^-16^). As a control, we further compared the mutation density within and out of MHC binding peptides in undetectably-expressed genes. Our results indicated that there was no significant depletion of missense somatic mutations within MHC binding peptides that are not detectably expressed (Figure 1; odds ratio = 1.01, *P*-value = 0.65). Although the odds ratio was near 1 for non-expressed proteins, as one might naively expect, we note that the sequence specificity of specific MHC class I receptor alleles can lead to HLA-allele-dependent amino acid (and therefore nucleotide-level) sequence biases in the peptides displayed, which could in turn yield sequence-dependent differences in mutation density. To account for this, we performed a correction by dividing the mutation density ratio of expressed proteins by that of non-expressed proteins. Although in this case the corrected mutational density ratio was 0.93/1.01, which is still 0.93, it did make a difference for other results below.

**Figure 1.**
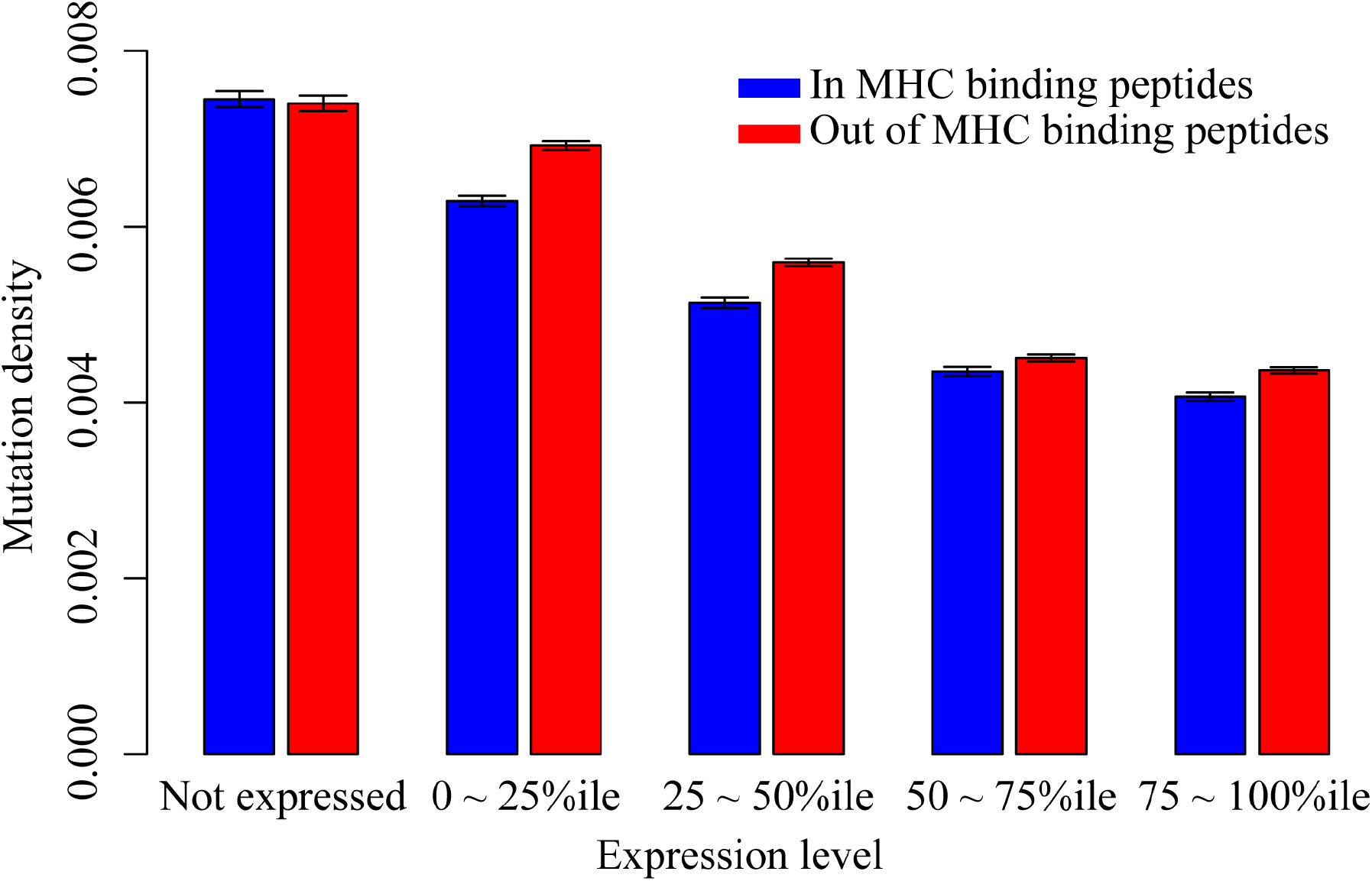
MHC-display-dependent mutation densities for genes with different expression levels. Blue bars are the mutation density within the predicted MHC binding peptides. Red bars are the mutation density out of the predicted MHC binding peptides. Mutations were separated into five categories based on the expression levels of their genes.

Thus, our analysis of PCAWG data confirmed the expected phenomenon that somatic mutations are depleted within expressed MHC-displayed peptides. Quantifying the MHC-display-dependent depletion effect in non-expressed peptides served as a crucial negative control for sequence biases of peptides displayed by particular HLA alleles.

### Depletion of mutations within predicted patient-displayed MHC-binding peptides

For a mutant protein to yield a peptide that is displayed by a given allele of the MHC class I receptor, that allele must of course be present in the cells of that patient. Because the analyses above were based on a hypothetical (and unrealistic) patient who bears all 12 of the common HLA alleles for which display predictions are available, the depletion effect sizes estimated above are likely to be conservatively small. Indeed, individual patients can differ dramatically in their immune systems, in part due to allelic variation in HLA genes. Therefore, we sought to characterize the mutation depletion phenomenon using, for each somatic variant, only peptide display predictions for the subset of HLA alleles carried by the patient in which that somatic variant was detected.

Re-examining the PCAWG data, there were 12,552 genes in which at least one variant was predicted to be neo-antigenic, e.g., presented by the MHC class I protein of the patient carrying this mutated gene. For these genes, we again examined the tendency for depletion of mutations within MHC binding peptides relative to non-MHC binding peptides, now taking patient HLA type into account. Within expressed proteins, the ratio of mutation density within predicted-displayed MHC binding peptides to that outside predicted-displayed peptides was 0.82 (Fisher’s exact test, *P*-value < 2.2e^-16^). Within non-expressed proteins, the corresponding ratio was 0.98 (Fisher’s exact test, P-value = 0.19), yielding a corrected mutational density ratio for expressed proteins of 0.83 (0.82/0.98).

Our analysis showed that missense mutations tend to be counter-selected within MHC binding peptides, both in an idealized patient with unknown HLA type, and when accounting for HLA type in each specific patient sample. In each case, the phenomenon depended on expression level of the gene encoding that peptide (Figure 2). In all subsequent analyses, we considered only peptides expressed according to RNA-Seq analysis of the appropriately-matched cancer type.

**Figure 2.**
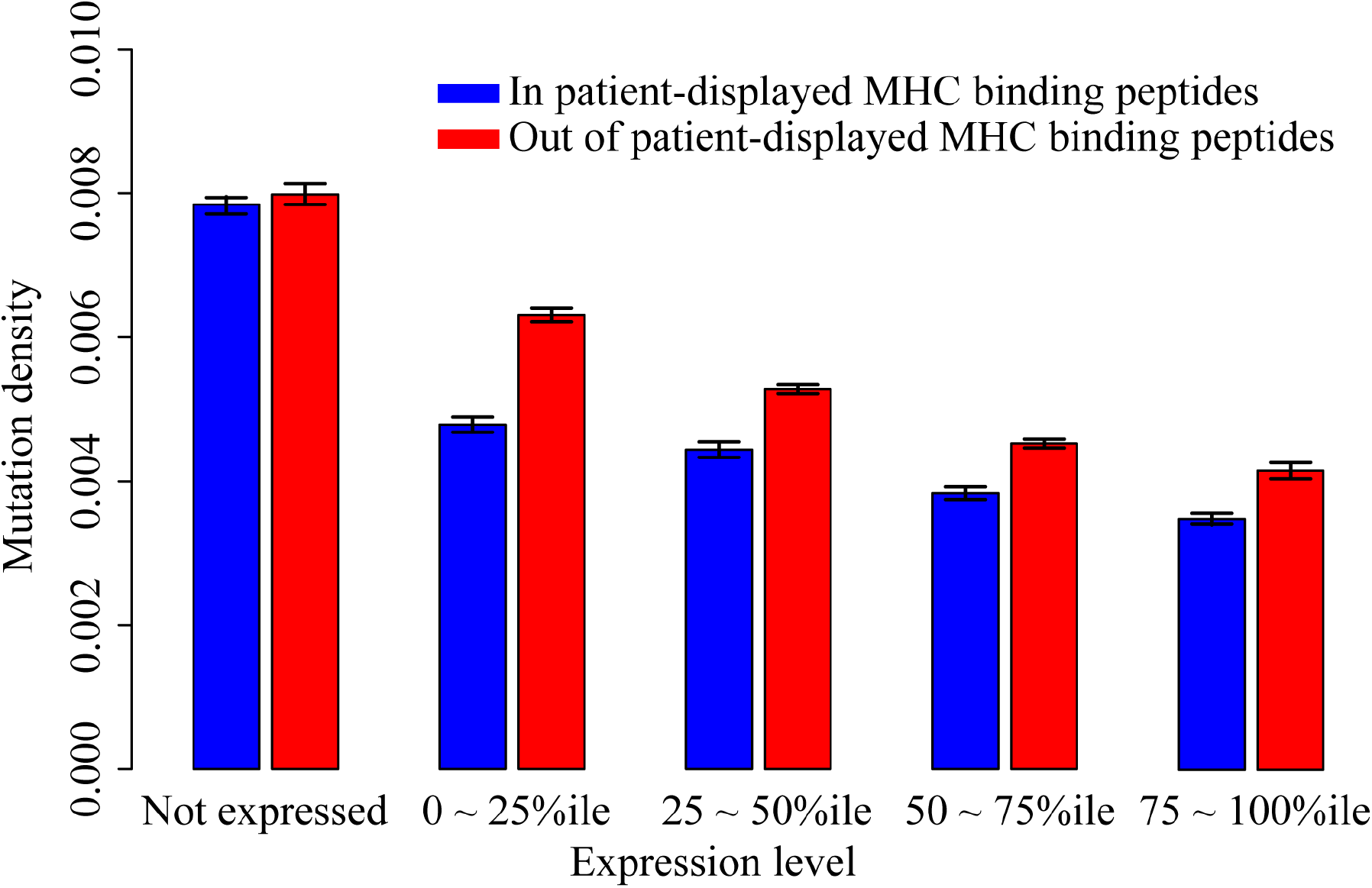
MHC-display-dependent mutation densities for genes with different expression levels, considering each patient’s HLA type. Blue bars are the mutation density within the predicted patient-displayed MHC binding peptides. Red bars are the mutation density out of the patient-displayed predicted MHC binding peptides. Mutations were separated into five categories based on the expression levels of their genes.

### Dependence of depletion on the number of mutation-displaying alleles

In the above analysis, we only considered for each peptide whether or not the patient carried an HLA allele predicted to display that peptide, but did not consider how many copies of the displaying allele were present in that patient. We hypothesized that peptides for which two copies of the displaying HLA alleles were present would be more efficiently displayed. (This could be due either to increased expression of the displaying allele by increased gene dosage, or a decreased chance that the displaying allele would be silenced where the phenomenon of mono-allelic expression occurs^13^). We assessed this hypothesis further by testing, for patient samples where ‘likely-displayed’ mutations were found, if the number of alleles that can display the MHC binding peptides was associated with the extent of mutation depletion.

Missense variants from the 2,834 PCAWG patient samples were separated into three types (Figure 3). “D0,” where the patient has zero HLA class I alleles that are predicted to display the mutant peptide; “D1”, where only one HLA class I allele type can display the peptide, i.e., the patient is heterozygous at the relevant HLA locus such that the patient has only one HLA allele that can display the peptide; and “D2”, where two HLA class I alleles are predicted to display the peptide.

**Figure 3.**
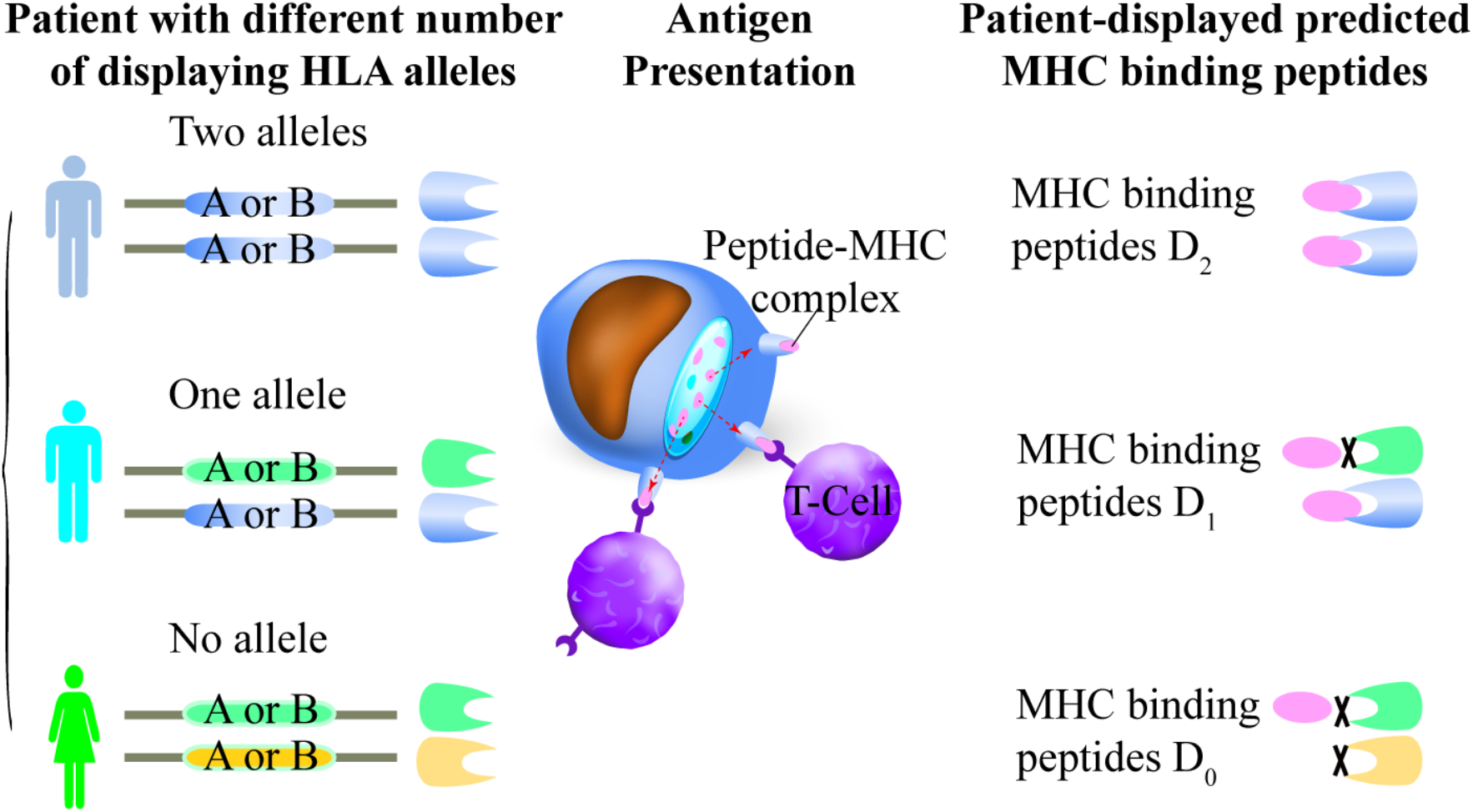
Three types of MHC binding peptides based on patient HLA allele types.

We found that for both D1 and D2 mutations, the mutation density within patient-displayed MHC binding peptides is lower than that observed outside of MHC binding peptides of the same protein. For expressed MHC binding peptides of type D1, the ratio of mutation density within displayed peptides to that outside of displayed peptides was 0.91 (Fisher’s exact test, *P*-value = 1.96e^-7^). This ratio for non-expressed peptides was 0.99 (Fisher’s exact test, *P*-value = 0.39), yielding a corrected mutational density ratio of 0.92 (0.91/0.99) for expressed D1 peptides.

For expressed displayed peptides of type D2, the ratio was 0.79 (Fisher’s exact test, *P*-value = 9.73e^-9^). The corresponding ratio in non-expressed displayed peptides D2 that can be displayed by two distinct HLA alleles is 1.02 (Fisher’s exact test, *P*-value =0.64). Thus, a corrected mutational density ratio 0.77 (0.79/1.02) was observed for expressed D2 peptides displayed by two HLA alleles.

Thus, we find that the depletion for mutations in MHC-displayed peptides is stronger if the patient has more alleles predicted to display a mutant peptide. Interestingly, our analysis showed that the depletion effect is stronger for mutants where the patient is homozygous for a single HLA display-enabling allele type than when the patient has two distinct HLA display-enabling alleles (Figure 4).

**Figure 4.**
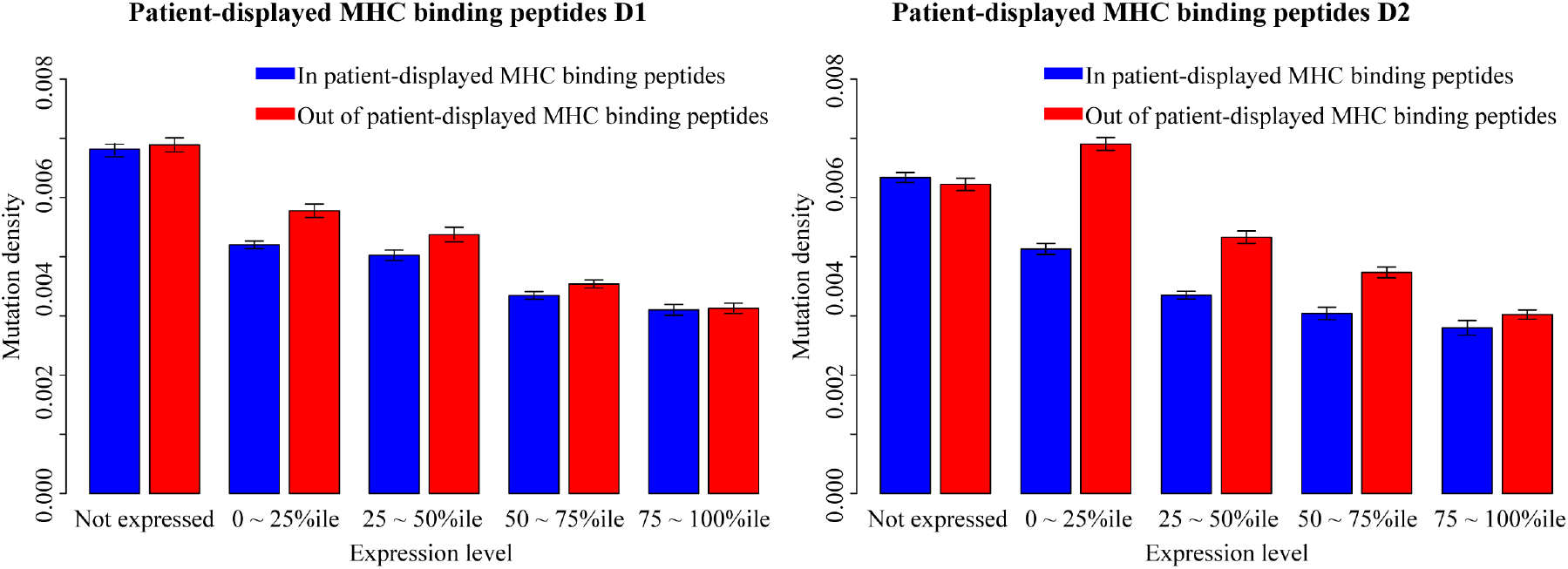
MHC-display-dependent mutation densities for genes with different expression levels, considering the number of displaying HLA alleles. Average mutation density in peptides predicted to be displayed by one or two of the 12 common HLA-A or HLA-B allele types. A. Mutation density in peptides predicted to be displayed in patients by only one HLA allele. B. Mutation density in peptides predicted to be displayed in patients with two displaying HLA alleles.

### Validation of mutation depletion phenomena in an independent dataset

We repeated the above analyses using the missense somatic mutations detected from 5,213 patient samples provided by the TCGA project^14^, examining the distribution pattern of 676,171 missense mutations detected in more than 10,800 genes. Analysis of this TCGA data confirmed the tendency of depletion of mutations within MHC binding peptides relative to non-MHC binding peptides, both with and without considering patient HLA types (Figure S1, S2). Considering only patient-displayed MHC binding peptides, the corrected mutational density ratio was 0.54 (with 95% confidence interval of 0.539 – 0.545 estimated by bootstrap resampling; Figure S3). Our analysis of the TCGA data confirmed that mutations displayed by two display-enabling HLA alleles (mutations of type “D2”) were more strongly depleted than mutations displayed by a single display-enabling allele (Figure S4).

To address concerns that the depletion phenomenon stems from a bias in the spectrum or rate of mutation for expressed genes, we also analyzed 1,048,575 synonymous mutations in 5,134 samples. We did not find depletion of synonymous mutations within patient-displayed MHC binding peptides (Figure S5). Within expressed proteins, the ratio of synonymous mutation density within predicted-displayed MHC binding peptides to that outside predicted-displayed peptides was 1.05 (Fisher’s exact test, P-value = 0.99). Within non-expressed proteins, the corresponding ratio was 1.03 (Fisher’s exact test, *P*-value = 0.78), yielding a corrected mutational density ratio for expressed proteins of 1.01 (1.05/1.03). The 95% confidence interval of the corrected mutational density ratio for synonymous variants was 1.00 to 1.01 (based on bootstrap resampling 500 times; Figure S6). That we observed no depletion of synonymous mutations in patient displayed MHC binding peptides is consistent with the hypothesis that the depletion phenomenon arises from a selection that depends on expression of the mutant protein.

We next repeated our analysis by considering different cancer types separately. Here, we chose the six different types for which the most samples were available: breast cancer (BRCA, 973 samples), thyroid cancer (THCA, 386 samples), skin cutaneous melanoma (SKCM, 341 samples), prostate adenocarcinoma (PRAD, 329 samples), gastric adenocarcinoma (STAD, 275 samples) and uterine corpus endometrial carcinoma (UCEC, 240 samples (see Table S2). Significant depletion of predicted-displayed mutations had (without considering patient HLA type or peptide expression) been previously found for BRCA and STAD^10^. We also included adenomatous colorectal cancer (COAD, 60 samples), because Rooney *et al*. noted highly-significant depletion for this cancer type. Considering patient HLA genotypes and proteins in the 75%-100% expression levels percentile range, we were able to confirm the trend of depletion of mutations in MHC-binding peptides for BRCA, STAD and COAD, and also THCA (not noted previously). Although we could not confirm depletion of mutations in UCEC, SKCM and PRAD at 75%-100% expression percentile, depletion was seen for all three of these at other expression percentiles (Figure S7).

As a negative control, we performed the same analysis for synonymous mutation density within predicted-displayed MHC binding peptides relative to that outside predicted-displayed peptides. This ratio did not vary significantly from unity for any of the seven cancer types: The ratio and Fisher’s exact test P-values for each cancer were: COAD: 0.97, 0.25; BRCA: 0.96, 0.04; THCA: 0.94, 0.26; STAD: 0.96, < 2.2e^-16^; UCEC: 0.99, 0.31; SKCM: 0.95, 5.9e^-5^; and PRAD: 1.00, 0.52. For non-expressed genes, the corresponding results were COAD: 1.04, 0.67; BRCA: 1.01, 0.56; THCA: 1.03, 0.60); STAD: 0.98, 0.42; UCEC: 1.02, 0.63; SKCM: 0.97, 0.09; and PRAD:1.00, 0.54. Only for BRCA were there enough samples to separate mutations into the three categories, D0, D1 and D2, although even for BRCA only 8-10 mutations fell into the D2 category (Figure S8). Although we could not find significant depletion of mutations within the patient displayed MHC binding peptides within each cancer type, this result indicate that the counter-selection of nonsynonymous mutations might exist in even more cancer types once we have more samples sequenced.

## DISCUSSION

In this study, we examined signatures of immune selection pressure on the distribution of somatic mutations, quantifying the extent to which somatic mutations are significantly depleted in peptides that are predicted to be displayed by MHC class I proteins, and characterizing the dependence of this depletion on expression level. We also examined whether immune selection pressure on somatic mutations changes depending on whether there are either one or two HLA alleles that can display the peptide.

Only expressed MHC binding peptides that can be displayed by at least one patient HLA allele are immunogenic in terms of class I MHC display. In our analysis using the PCAWG dataset, we found mutation densities to be similar for mutations within or out of the predicted MHC binding peptides when the gene was not expressed (Figure 1). That proteins must be expressed to be antigenic is one explanation for the fact that many “likely-displayed” mutations were nevertheless observed in a tumor. More refined estimates of the depletion effect in future studies might come from using expression data from a specific patient tumor sample.

We note that the terms “in MHC binding peptides” and “out of MHC binding peptides” were applied based on whether or not peptides were predicted to be displayed by at least one of the 12 common HLA-A or HLA-B allele types. We expect that the phenomenon of depletion of somatic mutations “out of MHC binding peptides” is observed where patients do not have a common displaying allele type because such peptides are more likely to be displayed by one of the HLA-C alleles or less common HLA-A or HLA-B allele types or patients with different specific HLA type combinations.

We expect that this information will be useful in building a model that predicts the antigenicity of any given missense mutation detected by whole genome or whole exome sequencing. Although scores for observed mutations based on counter-selection of similar mutations may over-estimate neoantigenicity (if a somatic mutation has been observed, it has obviously not yet been cleared by the immune system), such scores could point to ‘cryptic immunogenicity’ of a somatic variant. In cases of cryptic immunogenicity, some therapies might enable immune clearance of cancer cells by ‘revealing’ this immunogenicity, e.g. by de-silencing HLA loci within cancer cells, or by relieving tumour-derived suppression of immune cells. These results would therefore be potentially useful in scoring tumors with greatest potential to benefit from immunotherapy, or may aid in developing personalized cancer vaccines that introduce or stimulate immune cells to recognize specific predicted neo-antigens.

Our results also supported the idea that having two copies of the display-enabling allele is more effective for peptide display than having just one copy. This could result from a gene-dosage effect, or via monoallelic expression (MAE). MAE, the phenomenon that only one allele of a given gene is expressed, is a frequent genomic event in normal tissues. MAE-derived silencing of one or more HLA-encoded alleles could potentially cause failure to express MHC binding-peptide-encoding genes, which may, in turn, alter the immunogenicity of somatic mutations. A previous study showed that the genome-wide rate of MAE was higher in tumor cells than in normal tissues, and the MAE rate was increased with specific tumor grade. Oncogenes exhibited significantly higher MAE in high-grade compared with low-grade tumors^13,15,16^. The role of MAE in immunogenicity of cancerous cells is entirely unclear. Because HLA alleles are known to be subject to MAE^13^, it may be interesting in future studies to assess the impact of MAE by comparing the mutation rates between homozygous and heterozygous samples at HLA class I loci A and B respectively using the allele-specific expression data. One example of a potential therapy that might emerge from this study is that de-silencing (either global or targeted) could lead to the display of otherwise-cryptic neo-antigens and therefore to immune clearance of cancerous cells, especially when used in combination with current immunotherapy strategies. If we can better understand the interplay between individual immune systems and the likelihood that cancer cells bearing specific somatic mutations are cleared, we will gain insight into the therapeutic potential of MAE modulation. For example, if MAE can indeed limit peptide display efficiency, then therapies reducing MAE could potentially increase the efficiency of immune clearance of tumor cells.

With the analysis conducted here, we can begin to quantify the efficiency of immune clearance of somatically mutated cells. For example, for somatic mutations in proteins expressed in a given cancer type, the depletion ratios we observed were as low as 0.77 (for expressed peptides predicted to be displayed by an MHC receptor encoded by two copies of the same HLA allele). This result allows us to conservatively predict that cells bearing somatic mutations falling within DNA segments encoding such peptides are cleared roughly 33% of the time by the immune system at tumor stages that are earlier than those examined in PCAWG sequencing studies. Because any inaccuracy in estimating protein expression levels or peptide display would be expected to diminish our ability to detect the depletion phenomenon, this estimate of immune clearance rate is likely conservative.

## MATERIALS AND METHODS

### Obtaining catalogs of somatic variants in cancer samples

This study made use of two different collections of cancer-cell-derived somatic variants. First, we used data from The Pan-cancer Analysis of Whole Genomes (PCAWG, May 2016 version 1.1) project^17,18^, including 121,258 missense somatic cancer mutations in 10,745 genes detected from 2,834 patient samples. The number of patient samples in each cancer type is shown in Table S1.

Second, we examined data downloaded from The Cancer Genome Atlas (TCGA) project, obtaining 676,171 missense somatic cancer mutations in 18,106 genes detected from 5,213 patient samples (Table S2). We also examined 1048,575 synonymous mutations in 5134 samples as a control. Data were downloaded from Broad Institute TCGA Genome Data Analysis Center (2016-01-28).

### Mapping somatic variants to proteins

Protein sequences were downloaded using BioMart R package^19^ based on the Ensemble Protein IDs provided in PCAWG and TCGA datasets. Each missense mutation was mapped to the corresponding protein based on the position of the mutation with respect to a given protein (Figure 5). Also, we validated that the wild type residue given for the mutation was found at the corresponding position within the downloaded protein sequence.

**Figure 5.**
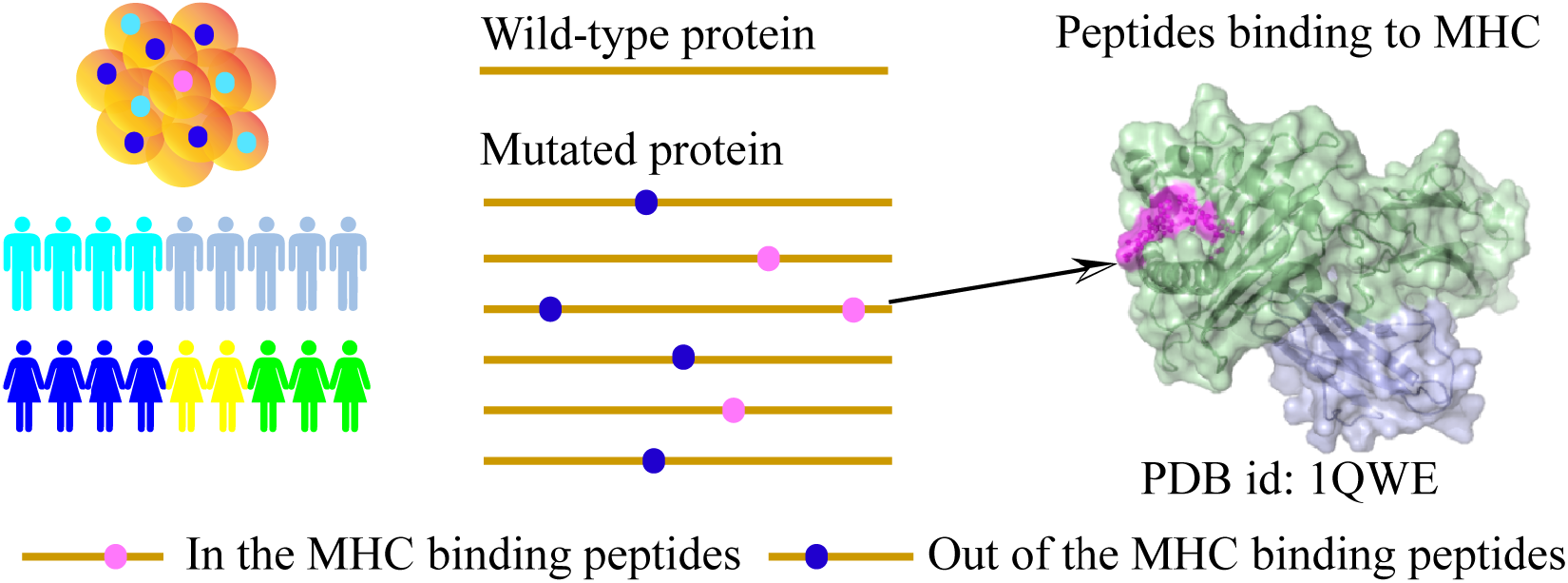
Predicting MHC-binding peptides and calculating mutation densities. Mutations within the MHC binding peptides are shown in blue dots, and mutations out of the MHC binding peptides are shown in pink dots. Protein sequence are shown as yellow line.

### Predicting peptides bound by class I MHC Receptors

We used the NetMHC server, version 3.4^11,12^ to predict MHC binding peptides associated with 12 common HLA class I alleles: HLA-A*0101, HLA-A*0201, HLA-A*0301, HLA-A*2402, HLA-A*2601, HLA-B*0702, HLA-B*0801, HLA-B*1501, HLA-B*2705, HLA-B*3901, HLA-B*4001, and HLA-B*5801. For this study, NetMHC scores were obtained for MHC binding peptides of length nine (Although it is possible for peptides with 10 or 11 residues to bind, this is less common and such cases are more difficult to predict). Also, only strong MHC class I binding peptides with NetMHC affinity score of 50 or less were selected (smaller NetMHC scores correspond to higher affinity).

### Calculating the depletion of mutations within MHC class I binding peptides

For each class of proteins and variants examined, we determined the total number of mutations falling within and outside of predicted MHC binding peptide regions for each protein. To test for significant differences in proportions of counts in different groups of peptides, we performed Fisher’s exact test using the “stats” package in R.

### Estimating transcript expression levels

We estimated gene expression levels for TCGA patient samples using TCGA RNAseq data^20^. Data were downloaded from Broad Institute TCGA Genome Data Analysis Center (2016-01-28). The expression level of each gene for each cancer type was estimated using the median expression level of that gene across all TCGA samples of that cancer type. Genes were classified as detectably expressed (RNA-Seq by Expectation Maximization (RSEM) normalized expression value > 0) or not (RSEM normalized expression value = 0). Detectably expressed genes were grouped into four expression quantiles according to the RSEM normalized expression value of expression level.

### Classifying human leukocyte antigen (HLA) types

For PCAWG samples, the four-digit HLA type for 2834 patients was determined (S.H., H.N. and S.I., unpublished method) and all HLA types are shown in File S1. For TCGA samples, the four-digit HLA type of the 5213 TCGA patients was predicted using PolySolver^14^.

### Data availability

PCAWG patient HLA types are provided in file S1. TCGA missense mutation data and TCGA RNAseq expression data are available via the Broad Institute TCGA Genome Data Analysis Center (2016) as “Analysis-ready standardized TCGA data from Broad GDAC Firehose 2016_01_28 run Broad Institute of MIT and Harvard (Dataset. https://doi.org/10.7908/C11G0KM9)”. Protein sequence data are publicly available via Ensembl BioMart (Release 89). TCGA patient HLA types are available via Broad Institute’s Firehose (Authorization Domain: TCGA-dbGaP-Authorized).

## Supplementary Figures

**Figure S1.**
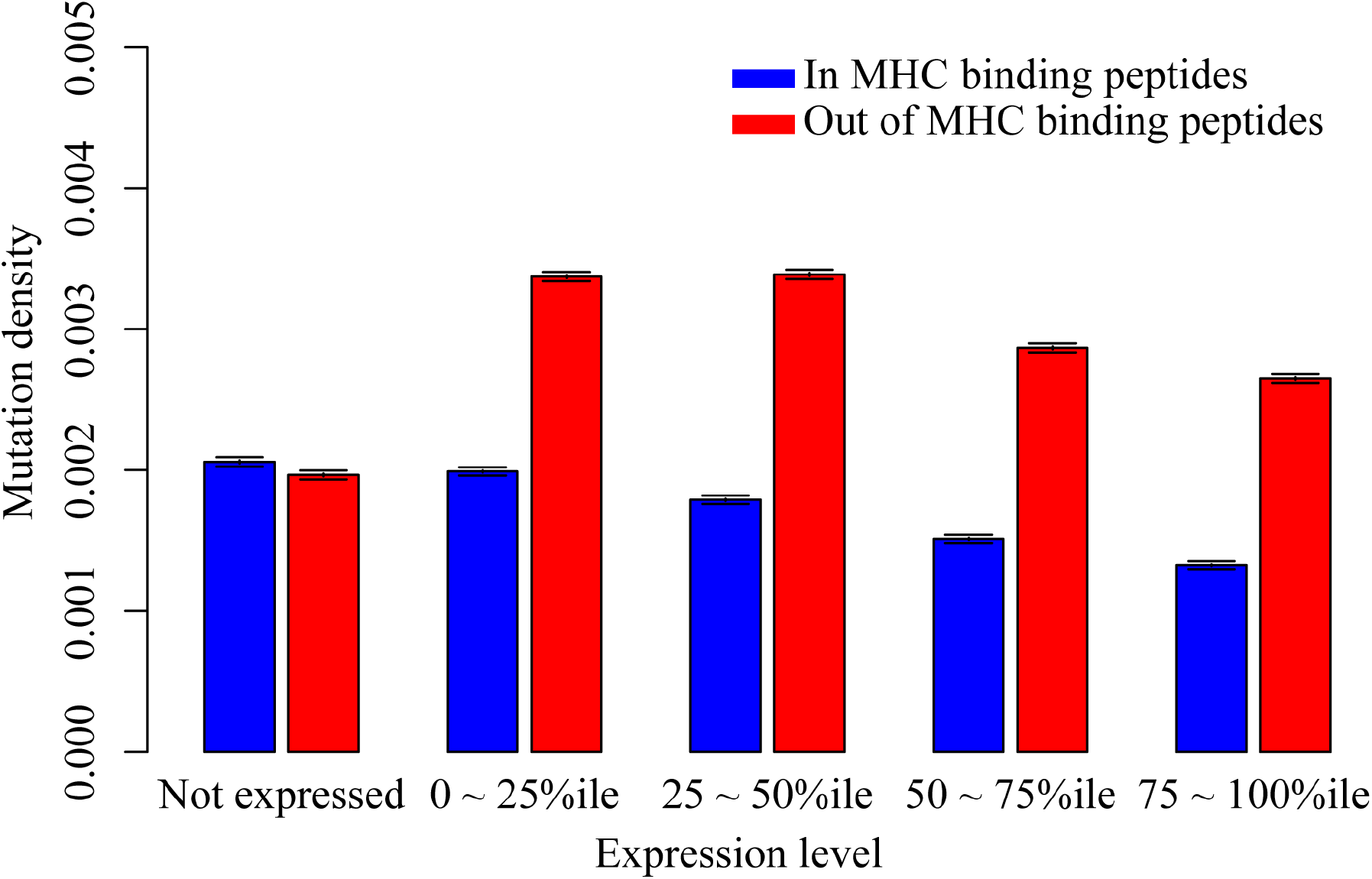
MHC-display-dependent mutation densities for genes with different expression levels using TCGA dataset. Blue bars are the mutation density within the predicted MHC binding peptides. Red bars are the mutation density out of the predicted MHC binding peptides. Mutations were separated into five categories based on the expression levels of their genes.

**Figure S2.**
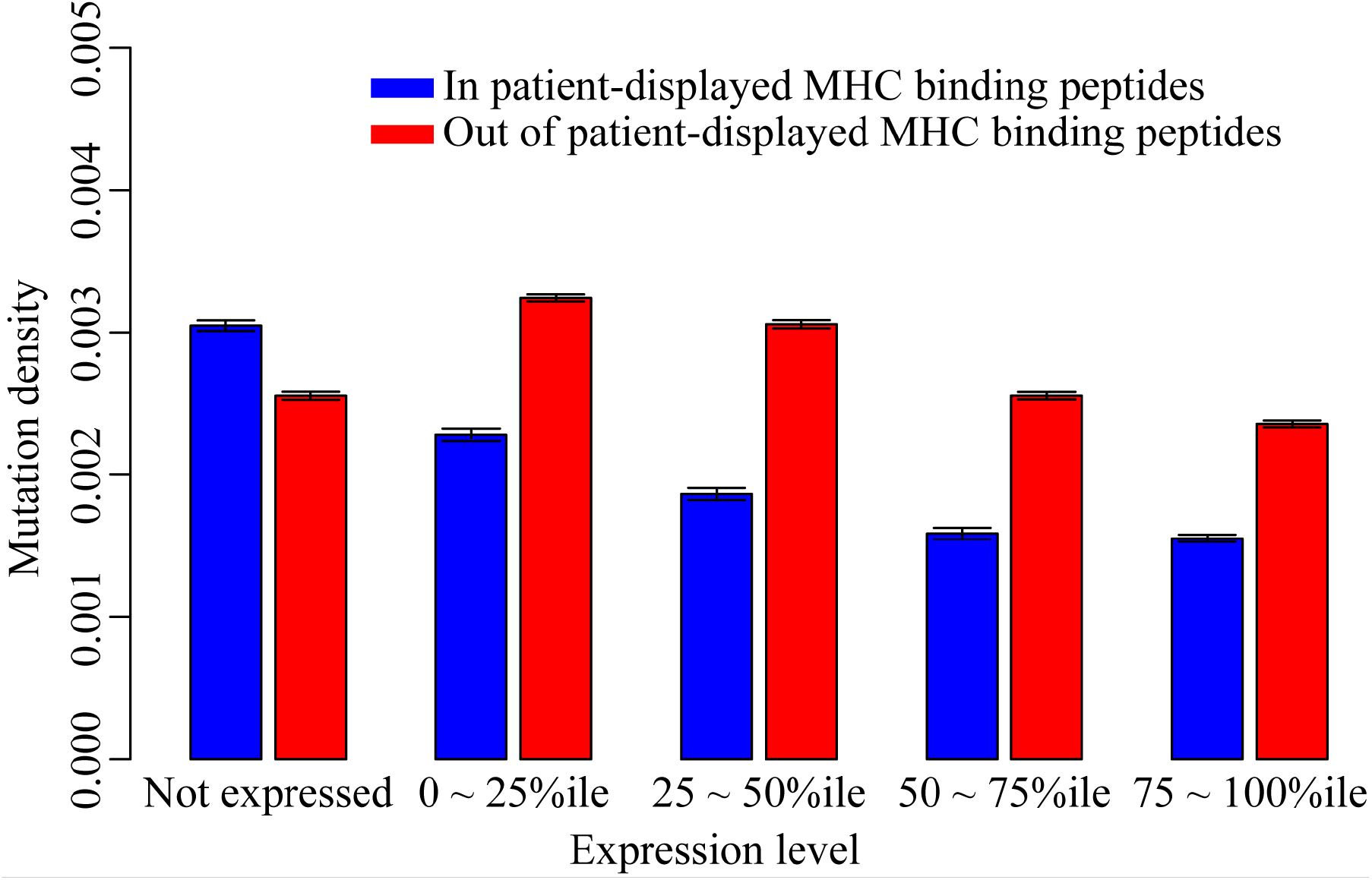
MHC-display-dependent mutation densities for genes with different expression levels, considering each TCGA patient’s HLA type. Blue bars are the mutation density within the predicted patient-displayed MHC binding peptides. Red bars are the mutation density out of the patient-displayed predicted MHC binding peptides. Mutations were separated into five categories based on the expression levels of their genes.

**Figure S3.**
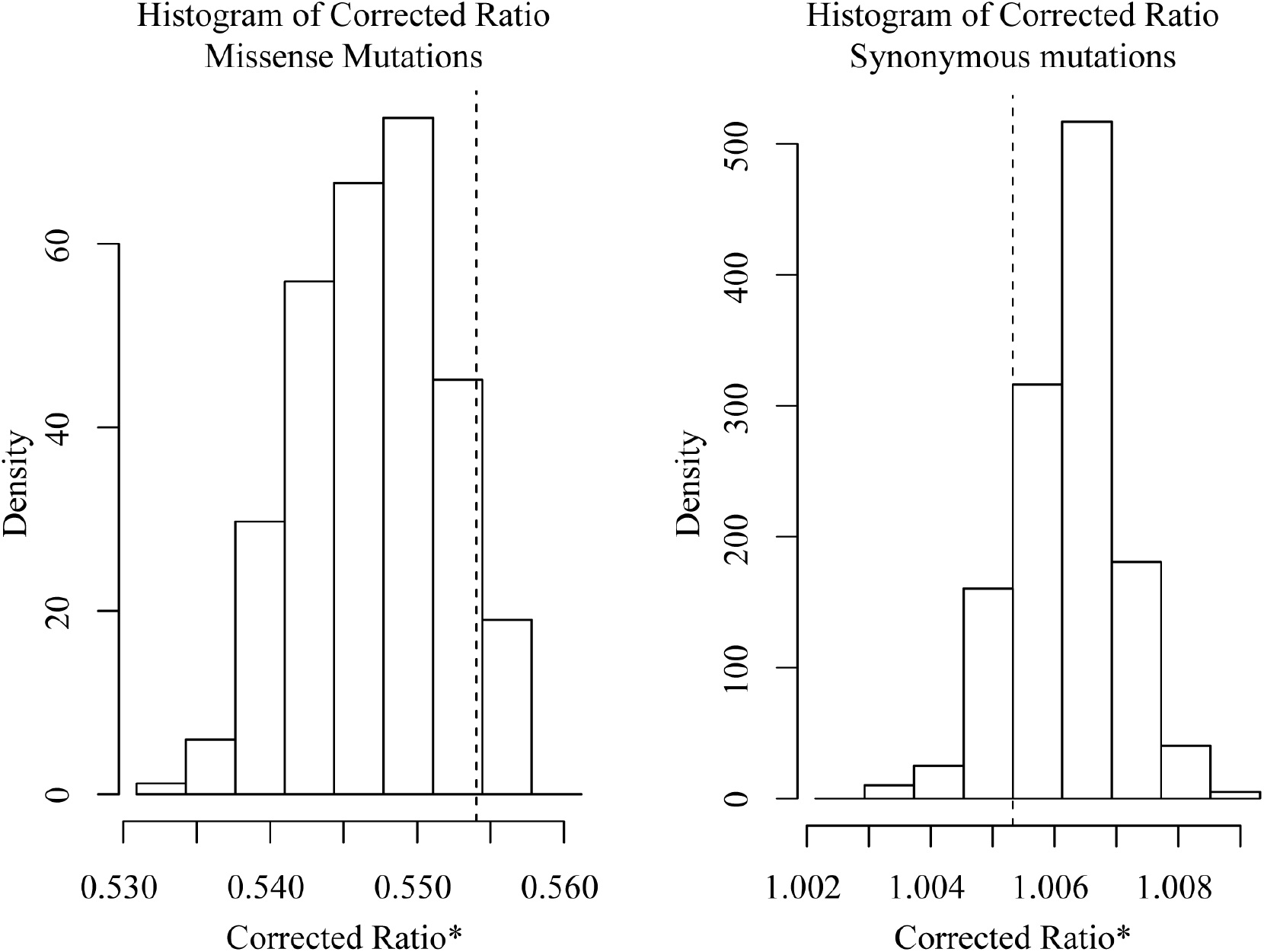
Exploring uncertainty in corrected mutation density ratio for TCGA mutations in patient-displayed MHC binding peptides, using bootstrap resampling, for both missense variants (left panel) and synonymous variants (right panel) Observed values are indicated with a vertical dashed line.

**Figure S4.**
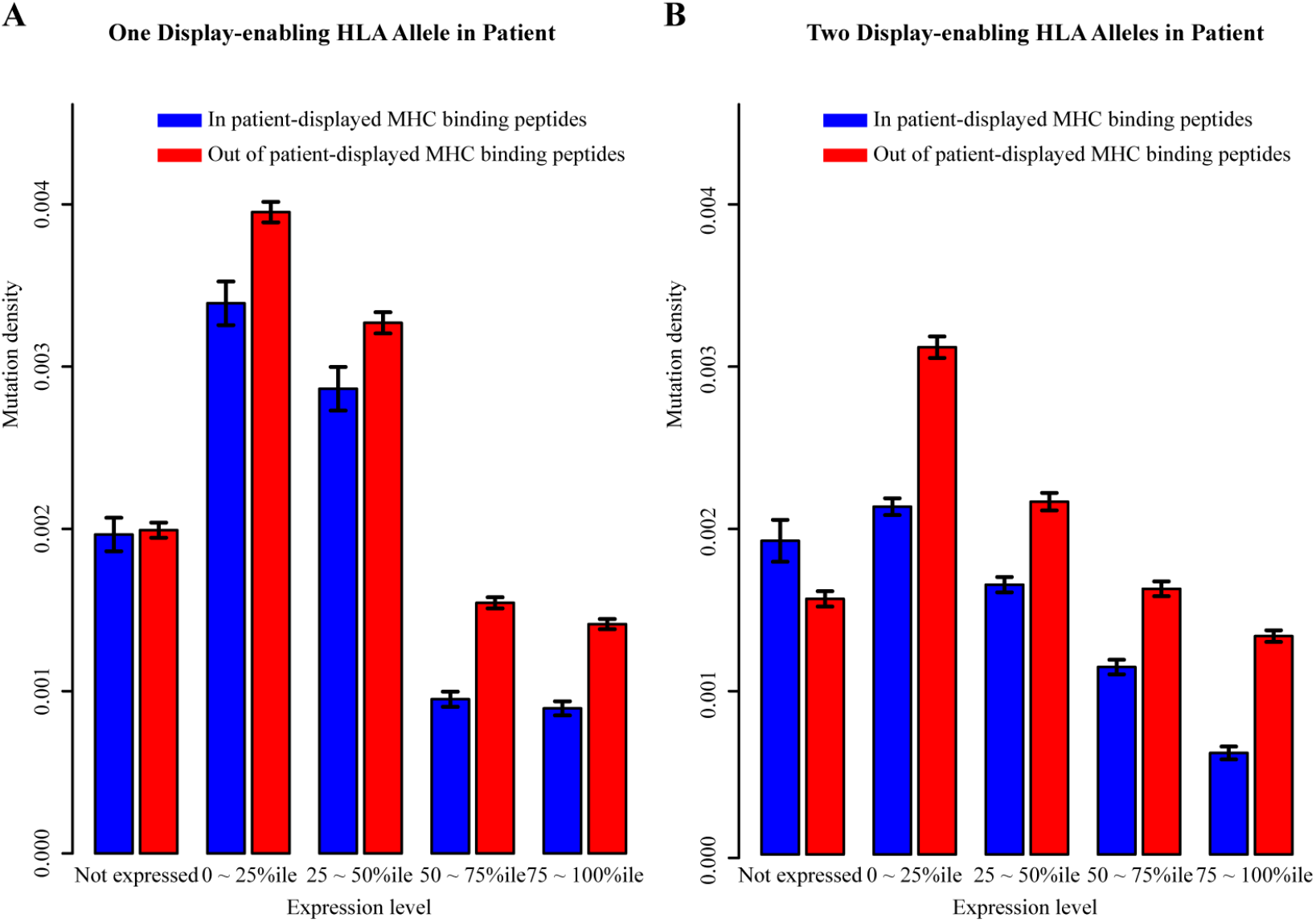
MHC-display-dependent mutation densities for genes with different expression levels, considering the number of displaying HLA alleles. Average mutation density in peptides predicted to be displayed by one or two of the 12 common HLA-A or HLA-B allele types. A. Mutation density in peptides predicted to be displayed in patients by only one HLA allele. B. Mutation density in peptides predicted to be displayed in patients with two displaying HLA alleles.

**Figure S5.**
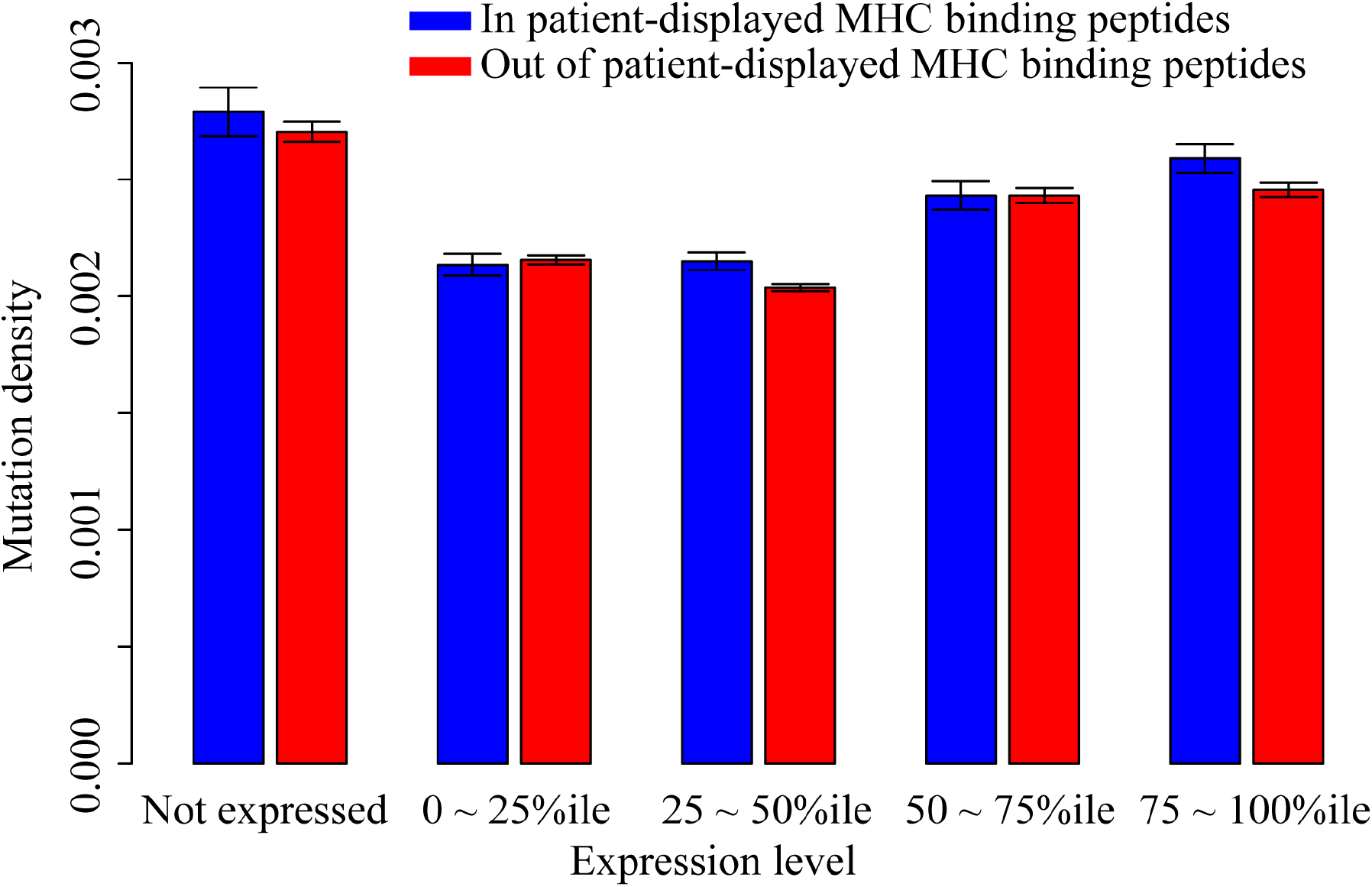
MHC-display-dependent synonymous mutation densities for genes with different expression levels, considering each TCGA patient’s HLA type. Blue bars are the synonymous mutation density within the predicted patient-displayed MHC binding peptides. Red bars are the synonymous mutation density out of the patient-displayed predicted MHC binding peptides. Synonymous mutations were separated into five categories based on the expression levels of their genes.

**Figure S6.**
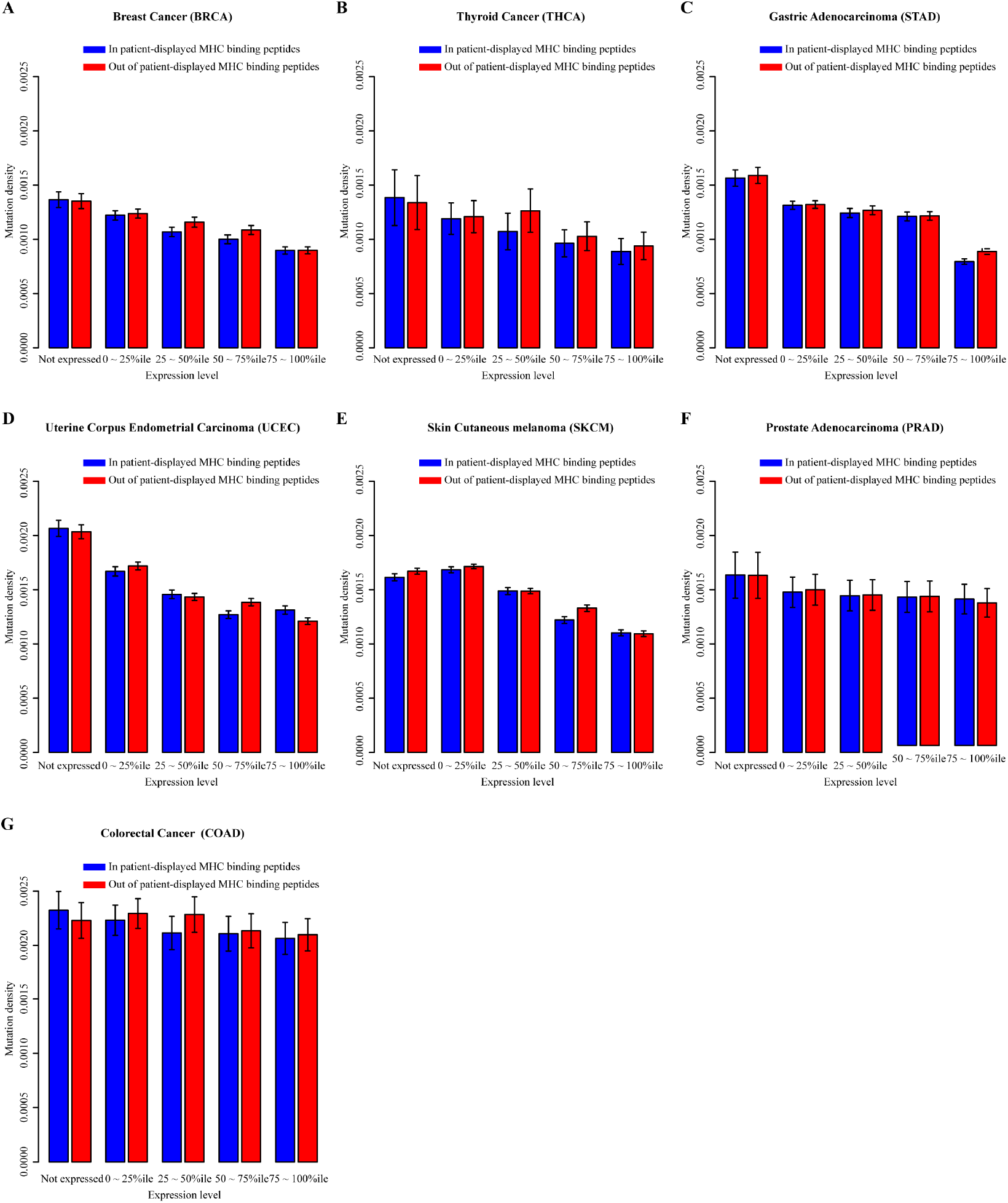
MHC-display-dependent mutation densities for genes with different expression levels in different cancer types. Blue bars are the mutation density within the predicted patient-displayed MHC binding peptides. Red bars are the mutation density out of the patient-displayed predicted MHC binding peptides. Mutations were separated into five categories based on the expression levels of their genes.

**Figure S7.**
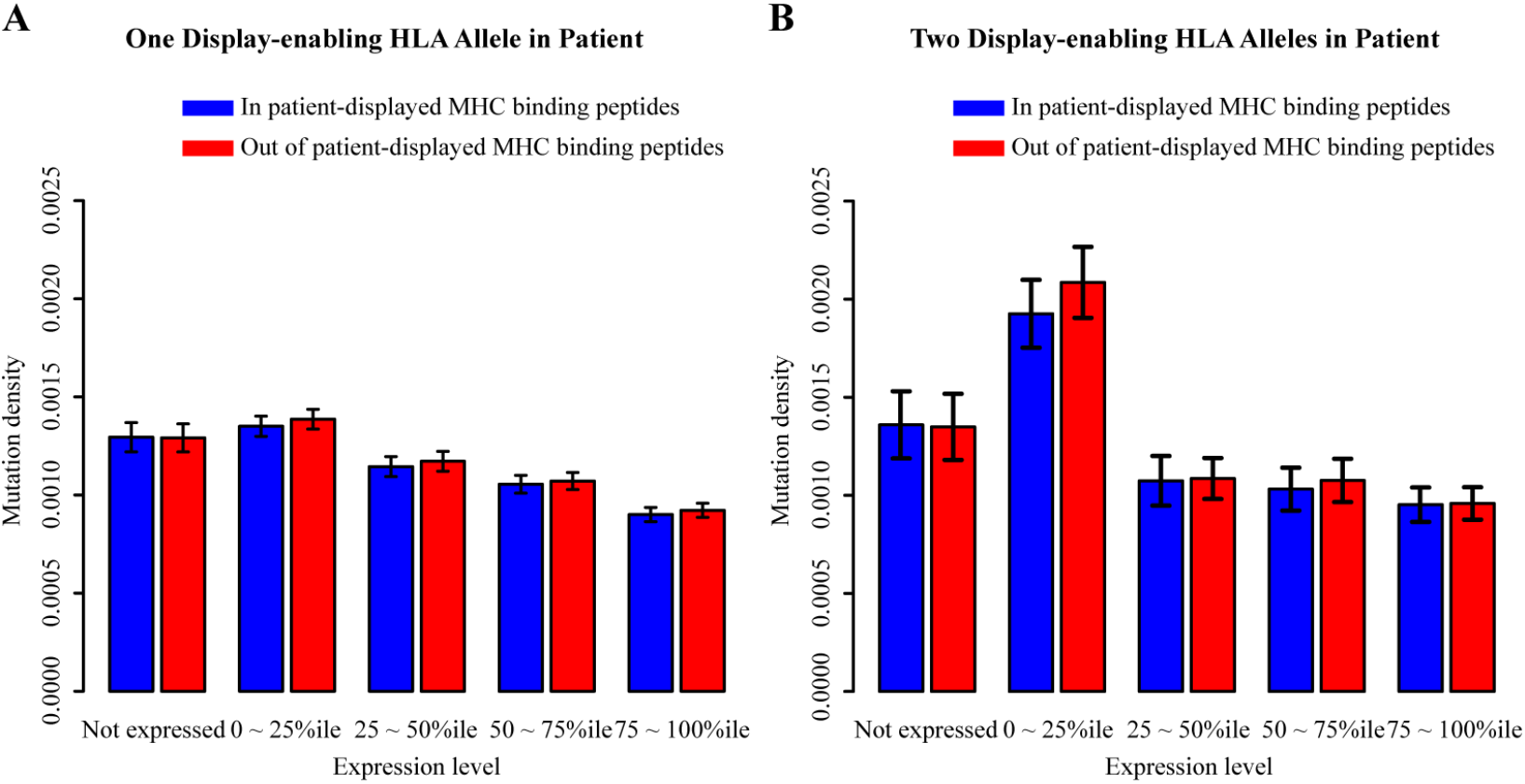
In Breast Cancer, MHC-display-dependent mutation densities for genes with different expression levels, considering the number of displaying HLA alleles. Average mutation density in peptides predicted to be displayed by one or two of the 12 common HLA-A or HLA-B allele types. A. Mutation density in peptides predicted to be displayed in patients by only one HLA allele. B. Mutation density in peptides predicted to be displayed in patients with two displaying HLA alleles.

## Supplementary Tables

**Table S1.** List of 37 Different PCAWG Cancer Types with Number of Samples and Mutated Genes of Each Cancer Type.

| **Cancer** | **Number of Samples** | **Number of Mutated Genes** |
| --- | --- | --- |
| BLCA | 23 | 4004 |
| BOCA | 61 | 1327 |
| BRCA | 207 | 10304 |
| BTCA | 11 | 719 |
| CESC | 20 | 956 |
| CLLE | 100 | 1293 |
| CMDI | 48 | 433 |
| COAD | 46 | 37666 |
| DLBC | 7 | 744 |
| EOPC | 41 | 771 |
| ESAD | 39 | 4787 |
| GACA | 27 | 2000 |
| GBM | 41 | 2604 |
| HNSC | 42 | 4899 |
| KICH | 49 | 1112 |
| KIRC | 40 | 2162 |
| KIRP | 34 | 1613 |
| LAML | 37 | 353 |
| LGG | 19 | 336 |
| LICA | 6 | 505 |
| LIHC | 52 | 3446 |
| LIRI | 39 | 2465 |
| LUAD | 40 | 7781 |
| LUSC | 48 | 10980 |
| MALY | 97 | 6479 |
| ORCA | 12 | 879 |
| OV | 114 | 6946 |
| PACA | 140 | 8675 |
| PAEN | 88 | 1846 |
| PBCA | 215 | 2057 |
| PRAD | 40 | 708 |
| READ | 16 | 14882 |
| SARC | 33 | 1179 |
| SKCM | 36 | 21278 |
| STAD | 39 | 10152 |
| THCA | 48 | 527 |
| UCEC | 50 | 19460 |

**Table S2.** List of 31 Different TCGA Cancer Types with Number of Samples and Mutated Genes of Each Cancer Type.

| **Cancer** | **Number of Samples** | **Number of Mutated Genes** |
| --- | --- | --- |
| ACC | 90 | 10805 |
| BLCA | 124 | 22991 |
| BRCA | 973 | 51806 |
| CESC | 149 | 21538 |
| CHOL | 35 | 3758 |
| COAD | 60 | 7305 |
| COADREAD | 60 | 7305 |
| DLBC | 14 | 1186 |
| ESCA | 181 | 32197 |
| GBM | 205 | 9340 |
| HNSC | 178 | 19679 |
| KIRP | 158 | 8861 |
| LAML | 109 | 815 |
| LIHC | 170 | 15863 |
| LUAD | 163 | 30987 |
| LUSC | 110 | 24512 |
| PAAD | 98 | 14334 |
| PCPG | 184 | 2128 |
| PRAD | 329 | 7723 |
| SARC | 238 | 10308 |
| SKCM | 341 | 161952 |
| STAD | 275 | 75745 |
| TGCT | 154 | 7598 |
| THCA | 386 | 4339 |
| THYM | 112 | 1553 |
| UCEC | 240 | 113697 |
| UCS | 7 | 7 |
| UVM | 8 | 8 |
| UCS | 57 | 6512 |
| UVM | 80 | 1319 |

